# Metabotropic NMDA Receptor Signaling Contributes to Sex Differences in Synaptic Plasticity and Episodic Memory

**DOI:** 10.1101/2024.01.26.577478

**Authors:** Aliza A. Le, Julie C. Lauterborn, Yousheng Jia, Conor D. Cox, Gary Lynch, Christine M. Gall

**Affiliations:** Departments of Anatomy and Neurobiology, University of California; Irvine, 92697, USA; Psychiatry and Human Behavior, University of California; Irvine, 92868, USA; Neurobiology and Behavior, University of California; Irvine, 92697, USA

**Author notes:** Corresponding Authors: C.M. Gall and G. Lynch, To whom correspondence should be addressed: Christine M. Gall, Communicating corresponding author Department of Anatomy and Neurobiology, University of California at Irvine 837 Health Sci. Road. Irvine, CA 92697-1275.

## Abstract

Men generally outperform women on encoding spatial components of episodic memory whereas the reverse holds for semantic elements. Here we show that female mice outperform males on tests for non-spatial aspects of episodic memory (“what”, “when”), suggesting that the human findings are influenced by neurobiological factors common to mammals. Analysis of hippocampal synaptic plasticity mechanisms and encoding revealed unprecedented, sex-specific contributions of non-classical metabotropic NMDA receptor (NMDAR) functions. While both sexes used non-ionic NMDAR signaling to trigger actin polymerization needed to consolidate long-term potentiation (LTP), NMDAR GluN2B subunit antagonism blocked these effects in males only and had the corresponding sex-specific effect on episodic memory. Conversely, blocking estrogen receptor alpha eliminated metabotropic stabilization of LTP and episodic memory in females only. The results show that sex differences in metabotropic signaling critical for enduring synaptic plasticity in hippocampus have significant consequences for encoding episodic memories.

## Introduction

Sex differences in learning were described in the 19^th^ century and have been a much discussed topic ever since^1-3^. Current analyses reflect a growing interest by psychologists and behavioral neuroscientists in episodic memory, a type of everyday encoding that includes the identities, locations, and temporal order of events^4,5^. Episodic memory organizes the flow of experience and in doing so is critical for diverse aspects of cognition including inferential thinking and imagination^6^. Although there are discrepancies in the literature^7,8^, men generally outperform women in spatial tests (e.g., three-dimensional mental rotations, navigation)^1,3,9,10^ whereas the reverse holds for verbal and non-spatial episodic memory tasks^1-3,11-13^. Women also outperform men in recalling stories^14^, a core episodic operation that places demands on brain systems that retrieve items in their correct order.

Whether, and to what extent, the above patterns reflect sex differences in biological substrates as opposed to education and social expectations^15,16^ is poorly understood. Evidence for a male advantage in spatial learning across several mammalian species strongly suggests that results in humans to some degree reflect evolutionarily conserved neurobiological mechanisms^17^. Comparable animal data are lacking for those aspects of episodic memory for which women outperform men. This is surprising given recent evidence that male rodents acquire ‘what’ and ‘when’ information along with spatial relationships (‘where’) in an episodic manner^5^, and that encoding in rodents, as in humans^18-20^, depends on hippocampus. In mice, males outperform females on encoding the ‘where’ element of an episode^21^, but it is not known if females have an advantage on non-spatial components.

Related to the above, there are sex differences in synaptic plasticity in the adult hippocampus^21-25^, and thus differences are likely for the neurobiological substrates of episodic memory. Both sexes require NMDAR-gated ion fluxes to induce memory-related long-term potentiation (LTP) but only females use local estrogen signaling to stabilize the potentiated state^22-24^. The possibility that males might rely upon non-ionic functions of the NMDARs rather than those of estrogen receptors for LTP consolidation has not been tested. Early evidence for such metabotropic (m-) NMDAR function involved demonstrations that use-dependent dephosphorylation^26,27^ and internalization of the receptors occur in the absence of channel opening. Malinow and colleagues^28-30^ then described multiple instances of NMDAR-driven effects in the presence of the channel blocker MK801. There is now evidence for m-NMDAR contributions^31^ to excitotoxic damage^32^ and glutamate-induced changes in spine size^33-35^. However, it is not known if these non-ionic processes are engaged by the naturalistic stimulation patterns commonly used to induce LTP, are critical for the enhancement of synaptic responses which defines potentiation, or contribute to the encoding of episodic memory.

The present studies investigated the above issues by first determining if there are sex differences in rodent encoding of the identity, location, and temporal order of cues using paradigms that do not include practice or overt rewards^5^. These experiments tested the proposition that the male advantage in episodic ‘where’ encoding are balanced by a female advantage in acquiring ‘what’ and/or ‘when’ information. We then evaluated the contributions and nature of metabotropic signaling supporting LTP in males and females with particular interest in activity-induced remodeling of the synaptic actin cytoskeleton. Results provide novel and striking evidence that in males m-NMDAR functions are critical for both LTP stabilization mechanisms and episodic memory, and that females substitute estrogen signaling for these m-NMDAR functions.

## Materials and Methods

### Animals

Experiments used 2-4 month old Sprague Dawley rats (Charles River) and 2-4 month old sighted-FVB129 mice of both sexes. Animals were group-housed (rat, 2-4/cage; mice, 3-5/cage) in rooms (68°F, 55% humidity) with 12-hr light/dark cycle (lights on 6:30AM), and food/water *ad libitum*. Mice were used for behavioral experiments as these tasks were previously validated in mice^5^. Rats were used for electrophysiological (except Fig. 3C) and immunolabeling experiments because their larger hippocampus allowed for precisely targeting specific laminae, and immunofluorescence and phalloidin paradigms were previously validated in rats^22^. Females were estrous-staged^21^ and used outside proestrus (estrus/diestrus). NMDAR subunit analysis used diestrus females. For electrophysiology and microscopy experiments, N denotes number of hippocampal slices from ≥3 animals (**Table S1**). Experiments were conducted in accordance with the National Institutes of Health Guide for the Care and Use for Laboratory Animals and protocols approved by the Institutional Animal Care and Use Committee at the University of California, Irvine.

### Behavioral Assays

To assess effects of sex and mNMDAR function on episodic memory, mice were tested in tasks that used multiple odor cues and did not involve repetition or reward^5,21,36,37^. Mice were handled for 2-min the day before experimentation. Serial ‘What’ and ‘When’ tasks used plexiglass arenas (30x25-cm floor, 21.5-cm height) and two jars; the simultaneous ‘What’ and ‘Where’ tasks employed larger arenas (60×60-cm floor, 30-cm height) and 4 jars as described^5,21^. Each jar contained a filter paper scented with an odorant dissolved in mineral oil (∼0.1 Pascals). The following odors, with letter identification, were used: (A) (+)-Limonene (≥97% purity, Sigma-Aldrich), (B) Cyclohexyl ethyl acetate (≥97%, International Flavors & Fragrances), (C) (+)-Citronellal (∼96% Alfa Aesar), (D) Octyl Aldehyde (∼99%, Acros Organics), (E) Anisole 99% (∼99% Acros Organics). These odors were previously confirmed to be saliently balanced in both sexes^21^.

### Serial ‘What’ task

A mouse was placed into an arena containing two unscented jars for 5-min and then lacking jars for 5 min. They were then exposed a series of three odor pairs (3-min each/5-min apart): A:A>B:B>C:C. Five-min following trial C:C, the mice were presented odorant pair A:D (familiar vs. novel odors, respectively) and allowed to explore for 3 min. A four-odor version of this task added an additional odor pair to the initial sequence (A:A>B:B>C:C>D:D) and used A:E for testing. This task was counter-balanced by using D as the first odor (D:D>B:B>C:C>A:A>D:E).

### Serial ‘When’ task

The task used the four-odor sampling sequence described above (A:A>B:B>C:C>D:D; 3-min each/5-min apart), and a final presentation of the first and second odors from the sequence (A vs. B; less vs. more recent) for testing.

### Simultaneous ‘What’ task

Mice were habituated to the arena containing four empty jars for 5 min. Jars were removed and, after 3-min, mice were exposed to four different odors (A:B:C:D) simultaneously for 5-min. At retention testing 48-hours later mice were reintroduced to the chamber containing three familiar (A:B:C) and one novel (E) odor.

### ‘Where’ task

Arena habituation and initial odor sampling was the same as for the simultaneous ‘What’ task. Three minutes after initial sampling, the odors were reintroduced with the position of two odorant jars from opposite corners (pair A:D or B:C) switched. The mice then freely explored the chamber for 5 min.

For drug studies, the initial odorant sampling was extended to 10-min to allow both sexes to learn. The 5-min retention trial was conducted 24-hrs later.

### Behavioral scoring

Sessions were digitally recorded and scored by an observer blind to group. Cue sampling time (t) was collected as the number of seconds the mouse’s nose was actively pointed towards the odor hole (∼0.5-cm radius). Calculations for the Discrimination Index (DI) across the tasks were as follows: ‘Where’ DI = 100 x (t_sum of switched pair_ − t_sum of stationary pair_)/(t_total sampling_); serial ‘What’ and ‘When’ DI = 100 × (t_novel_ − t_familiar_)/(t_total sampling_); simultaneous ‘What’ DI = 100 × (t_novel_ − t_mean familiars_)/(t_total sampling_). Z-score calculations were as follows: (mean DI_female_ – mean DI_male_)/(standard deviation_male_).

### Field Electrophysiology

Hippocampal slices were prepared using a McIllwain chopper (370-µm; transverse) and immediately transferred to an interface recording chamber with perfusion of oxygenated artificial cerebrospinal fluid (aCSF; 60-70 mL/hr, 31±1°C, 95% O_2_/5% CO_2_) that included (in mM): 124 NaCl, 26 NaHCO_3_, 3 KCl, 1.25 KH_2_PO_4_, 2.5 CaCl_2_, 1.5 MgSO_4_, and 10 dextrose (pH 7.4). Experiments were initiated 2 h later. Field excitatory postsynaptic potentials (fEPSPs) were elicited using a twisted nichrome wire stimulating electrode in CA1a or CA1c stratum radiatum (SR) and recorded with a glass pipette electrode (filled with 2M NaCl; R=2-3MΩ) in CA1b SR. Single-pulse baseline stimulation was applied with fEPSP amplitude at ∼40-50% of maximum population-spike free amplitude. Responses were digitized at 20kHz using an AC amplifier (A-M Systems, Model 1700) and recorded using NAC2.0 Neurodata Acquisition System (Theta Burst Corporation). LTP was induced using 10 burst train of theta burst stimulation (TBS: four pulses at 100Hz per burst, 200ms between bursts). For LTP-threshold analysis, TBS triplets were applied four times at 90 sec intervals^21,22^. Drugs were infused into the bath 1-3 hr before TBS.

### Whole-Cell Current-Clamp

Hippocampal slices (350-µm, transverse) from 8-week old male mice were prepared using a Leica Vibroslicer (VT1000S) and placed in a submerged recording chamber with constant oxygenated aCSF perfusion (2ml/min) at 32°C. Whole-cell recordings (Axopatch 200A amplifier, Molecular Devices) used 4–7 MΩ glass pipettes filled with (in mM): 140 CsMeSO_3_, 8 CsCl, 10 HEPES, 0.2 EGTA, 2 QX-314, 2 Mg-ATP, 0.3 Na-GTP. Bipolar stimulating electrodes were placed in the CA1 SR, 100-150 μm from the recorded cell. EPSCs were recorded with the holding potential at +40 mV for NMDAR amplitude (at 50ms from stimulation artifact) in the presence of 50 µM picrotoxin.

### Fluorescence Deconvolution Tomography (FDT)

For measures of basal synaptic protein levels, hippocampal slices (370-µm) were immersed in cold 4% paraformaldehyde (PFA) overnight. For LTP experiments, electrodes were placed in CA1a and CA1c SR for stimulation and CA1b SR for recording, all equidistant from the cell layer. After ∼5 min of stable baseline, one TBS train was applied to each polarity of each stimulating electrode (pulses at 2x baseline duration). Control slices continuously received 3/min pulses. Slices were harvested after a specified time post-TBS (3 min for pERK^22^, 7 min for pSrc^38^, and 15 min for pCAMKII^39^) and fixed overnight. Slices were sub-sectioned (20μm) and 6-8 sections from the top (interface plane) of each slice were slide-mounted and processed for dual immunofluorescence^22^.

The following primary antibodies (concentration; vendor, catalogue number, RRID) were used: goat anti-PSD95 (1:1500; Abcam, ab12093, AB_298846) with either rabbit anti-pCaMKII (Thr286/Thr287) (1:500; Upstate (now Millipore), 06-881, RRID:AB_310282) or rabbit anti-pERK1/2 (Thr202/Tyr204) (1:500; Cell Signaling 4377, AB_331775); Mouse anti-PSD95 (1:1000; Invitrogen, MA1-045, AB_325399) with rabbit anti-pSrc (Tyr419) (1:250; Invitrogen, 44-660G, AB_2533714); Rabbit anti-GluN1 (extracellular) (1:1000; Alamones Labs, AGC-001, AB_2040023), anti-GluN2A (1:500, Alamones Labs, AGC-002, AB_2040025), anti-GluN2B (1:500, Alamones Labs, AGC-003, AB_2040028), or anti-GluN2B Tyr1472 (1:300; PhosphoSolutions, P1516-1472, AB_2492182) with goat anti-PSD95 (1:1500, abcam ab12093; AB_298846). Secondary antibodies (all at 1:1000) included AlexaFluor donkey anti-goat 488 (Invitrogen, A32814, AB_2762838), donkey anti-rabbit 594 (A32754, Invitrogen, AB_2762827), donkey anti-mouse 594 (A21203, AB_141633), and donkey anti-rabbit 488 (A21206, AB_2535792).

FDT analyses were as described^22,40-42^. Image z-stacks (136x105x2-μm, 200-nm steps; 63X capture) were collected from the CA1 SR from ≥5-7 sections per slice and processed for iterative deconvolution (99% confidence; Volocity 4.0, PerkinElmer). Three dimensional (3-D) montages of each z-stack were analyzed for synaptic labeling using in-house software (c99, Java (OpenJDK IcedTea 6.1.12.6), Matlab R2019b, PuTTY 0.74, and Perl 5.30.0). For each image, labeling was normalized and thresholded, and erosion and dilation filtering was used to fill holes and remove background pixels to detect edges of both faintly and densely labeled structures. Objects were then segmented based on connected pixels above a threshold across each channel separately. All immunofluorescent elements meeting size constraints of synapses and detected across multiple intensity thresholds were quantified. PSD95-immunoreactive elements were considered double-labeled with the second antigen if there was contact or overlap in fields occupied by the two fluorophores as assessed in 3-D. Approximately 20-30 thousand synapses were thus analyzed per z-stack. Based on the maximum intensity of each image, counts of double-labeled puncta were assigned to ascending density (fluorescence intensity) bins and the data were expressed as frequency distributions. Labeled puncta with immunofluorescence density at ≥95 were considered densely-labeled. Counts of densely-labeled puncta peer section were averaged with those from other sections of that slice to generate the mean slice value presented.

### F-actin phalloidin immunolabelling

AlexaFluor 568-conjugated phalloidin (Invitrogen; A12380) was diluted in water to 12 µM stock and then to 6 µM in aCSF (1% DMSO) prior to experimentation. Electrode placement and stimulation was as for FDT analyses. Beginning 3 min post-stimulation, phalloidin (6 µM, 2 µl) was applied topically onto the slice (3 times, 3-min apart)^43^. Three minutes after the last application, slices were fixed in cold 4% PFA overnight. After cryoprotection (20% sucrose in 4% PFA), slices were subsectioned, slide-mounted, washed in 0.1 M PB (10 min) and cover-slipped with Vectashield using DAPI (Vector Labs). To quantify spine phalloidin labeling, image z-stacks were captured as for FDT. Every image of each z-stack then received a small saturated 1x1-µm reference square to two corners of the image (Python 3.0). The global reference square adds a fixed maximum intensity level for all images without significantly altering the background or raw intensity values of phalloidin-labeled puncta; this step was added because the software assigns the final density values for phalloidin labeling based on the maximum intensity a given image. The image z-stacks were then processed for quantification of spine-sized puncta as described for FDT; labeled puncta within the density bins of ≥90 were considered to have dense concentrations of F-actin. Counts of densely-labeled puncta were then averaged across tissue sections to generate a mean value per slice. Values from experimental groups were normalized to those of their respective control group.

### Drug Administration

For behavior, Ro25-6981 (Ro25; 5mg/kg, saline) and methyl-piperidino-pyrazole (MPP; 0.6 mg/kg, 2% DMSO/saline) were injected intraperitoneally 30 and 60 min before exposure to odors, respectively. For electrophysiology, compounds were introduced to the slice bath via a syringe pump (6ml/hr) into the aCSF infusion line for final bath concentrations: MK801 (30µM; Tocris, 0924), APV (100µM; Hello Bio, HB0225), DNQX (20µM; Hello Bio, HB0261), picrotoxin (30µM; Sigma-Aldrich, P1675) in water. MPP (3µM; Tocris, 1991) and Ro25-6981 (3µM; Hello Bio, HB0554) were dissolved with DMSO (≤0.01%).

### Statistics

Data are presented as mean ± s.e.m values; statistical analyses (p-values, degrees of freedom, and t-tests) are presented in **Table S1**. Significance (p<0.05) was determined using GraphPad Prism (v6.0). For electrophysiology, the magnitudes of LTP (averaged fEPSP slopes for last 5 min of recordings, normalized to 20-min baseline) and STP (averaged over 1-min post-TBS) were compared via two-tailed unpaired Student’s t-test. TBS area analysis and STP (for threshold TBS) were analyzed with repeated-measures two-way ANOVA. For imaging and behavioral studies, two-tailed unpaired Student’s t-test was employed for comparing two groups. For ≥3 group comparisons, one-way ANOVA (post-hoc Tukey test) and two-way ANOVA (post-hoc Tukey) were used.

## Results

### Sex differences in episodic learning

People organize memory for the flow of everyday experience into discrete episodes that minimally contain information about events, locations, and sequences. Episodic encoding occurs without rehearsal or reinforcement and, in these and other ways, is distinct from operant learning typically used in animal experiments^4^. We used olfactory cues to evaluate episodic memory in mice^5^ because odors are of innate interest to macrosmatic animals. To assess ‘what’ encoding, mice were exposed to a sequence of three different odor pairs (A>B>C), followed by a retention trial that paired a previously exposed odor with a novel odor (A vs. D). As rodents preferentially investigate novel stimuli, more time spent exploring the novel vs. previously sampled cue indicates that the latter was remembered^5,36^. Both sexes preferred the novel odor and had similar retention scores (discrimination indices [DI] for males and females: 40.9±7.2 and 41.8±7.8, respectively; p=0.94, unpaired t-test) (**Fig. 1a**). When presented with four odor pairs in sequence, females again exhibited high retention scores whereas males did not (male vs. female DI: 7.3±4.2 vs. 35.3±4.5; p=0.0003, **Fig. 1a**). These results constitute evidence for a female advantage in acquiring a fundamental component of episodic memory (i.e., cue identify). We reevaluated this point using a version of the episodic ‘what’ task in which mice were allowed to freely investigate four different odors (A-B-C-D) for 5 minutes. At testing 48 hours later, one of the cues was replaced with a novel odor. Females recognized the replacement odor, but males did not (**Fig. 1b**).

**Fig. 1.**
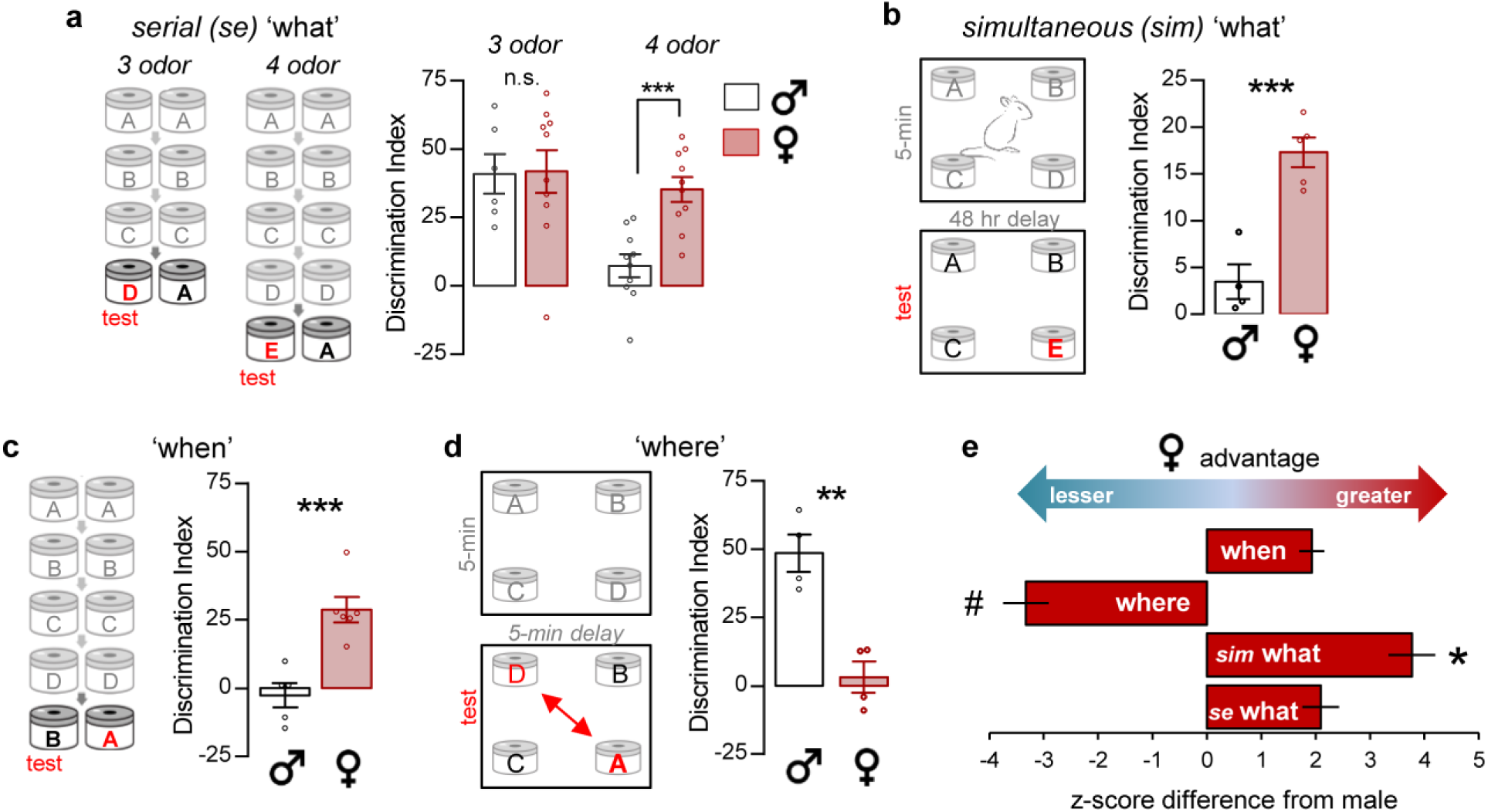
Females outperform males in tests for episodic ‘what’ and ‘when’ encoding. (**a**) Left: Schematic of the serial 3-odor and 4-odor ‘what’ tasks. Right: In the 3-odor task, mice preferentially explored the novel (D) vs. familiar (A) odor at testing with no sex difference (p=0.94; male N=6, female N=10). The presence of four odors in the test sequence severely degraded performance in males but not females (p=0.0003; male vs. female, N=10/group). (**b**) Simultaneous ‘what’ task schematic. Females, but not males, distinguished the novel from previously experienced odors (p=0.007; male N=4, female N=5). (**c**) In the ‘when’ task, females discriminated the least recently sampled odor whereas males did not (p=0.0021; male N=5, female N=6). (**d**) ‘Where’ task schematic. Males preferentially explored the novel-location odors whereas females did not (p=0.0022, N=4/group). (**e**) Female performance expressed as a z-score difference from the male group mean. The female advantage for simultaneous ‘what’ was greater than for the other tasks (F_3,21_=49.11, p<0.0001; *p≤0.15 Tukey post-hoc; N=4-10/group) and ‘where’ differed from the other three scores (# p<0.0001). Statistics: (**a-d**) two-tailed unpaired t-test; (**e**) One-way ANOVA, post-hoc Tukey. n.s. = not significant, *p<0.05, **p<0.01, ***p<0.001. Mean±s.e.m. values shown. **Table S1** contains detailed statistics.

Next, we tested for sex differences in encoding the temporal order of cue presentation (episodic ‘when’). Previous studies showed that male mice exposed to four consecutive odor pairs in series (A>B>C>D) spent more time investigating the less recent odor B (vs. more recent odor C) in retention testing^5^. This result obtained when the initial odor presentations were separated by 30 sec or 5 min, suggesting that mice acquire information about the order of cue presentation as opposed to the time since last exposure. Here, we used the same initial odor exposures but in the retention trial placed a heavier demand on memory by comparing the temporally more distant cues A vs. B. In contrast to results for B vs. C, males had no evident preference in A vs. B trials whereas females showed a clear preference for A over B and thus outperformed males in this regard (**Fig. 1c**). Finally, we tested spatial encoding (episodic ‘where’) by allowing mice to sample four simultaneously presented odors for 5 min and then tested if they detect cues shifted to novel positions. Males preferentially explored the switched (novel location) odors whereas females did not (**Fig. 1d**).

We summarized the results for the four episodic memory tests by expressing retention for each female mouse as a z-score difference from the mean of the male group. This provided a relative advantage-disadvantage estimate for female performance in each assay. The main effect was highly significant (F_3,21_=49.11, p<0.0001) with the strongest female advantage being in the simultaneous ‘what’ test (p<0.015 vs. other tests) and a marked female disadvantage in the ‘where’ test (p<0.0001 vs. other tests) (**Fig. 1e**). It is noteworthy that the same initial sampling trial used in both the simultaneous ‘what’ and ‘where’ tasks, yielded the greatest sex differences depending on which aspect of learning – cue identity vs. spatial location – was tested.

There were no systematic, cross-paradigm sex differences in the time spent sampling cues during initial sampling or retention trials. Similarly, travel distance and velocity were comparable between sexes (**Fig. S1**).

### Males use m-NMDAR signaling to consolidate LTP

#### Blocking the NMDAR channel does not interfere with stimulation-induced actin polymerization

Theta burst stimulation (TBS) of the CA3-CA1 projections causes a rapid and lasting increase in the density of filamentous (F-) actin in dendritic spines^44,45^ and blocking this effect prevents the stabilization of CA1 LTP^44,46-48^. To test if activity-driven actin polymerization requires NMDAR-mediated calcium influx we infused MK801, which occludes the NMDAR channel without interfering with glutamate binding to the receptor, prior to TBS. As expected, MK801 (30µM) produced a near complete suppression of both short-term potentiation (STP) and LTP (**Fig. 2a**). To evaluate effects on actin polymerization, we applied TBS to two populations of CA3 efferents converging on the apical dendrites of CA1b pyramidal neurons, with a 30-sec delay between activation of the inputs (**Fig. 2b**). AlexaFluor 568-phalloidin, which selectively binds F-actin, was topically applied after TBS and numbers of densely phalloidin-labeled spines in the CA1 apical dendritic sample field were counted (**Fig. 2c,d**) as described^49-51^. TBS robustly increased the number of spines containing dense phalloidin-labeled F-actin, and this effect was abolished by the competitive NMDAR antagonist APV. Remarkably, MK801, at the dose that eliminated LTP, did not attenuate the TBS-induced F-actin increase (**Fig. 2d**), indicating that activity-induced actin polymerization requires NMDARs but not the calcium influx mediated by those receptors. These results constitute evidence that naturalistic patterns of afferent activity initiate actin regulatory m-NMDAR signaling in adult synapses and describe a surprising instance in which a late-stage LTP stabilization event (actin polymerization) occurs in the absence of synaptic potentiation.

**Fig. 2.**
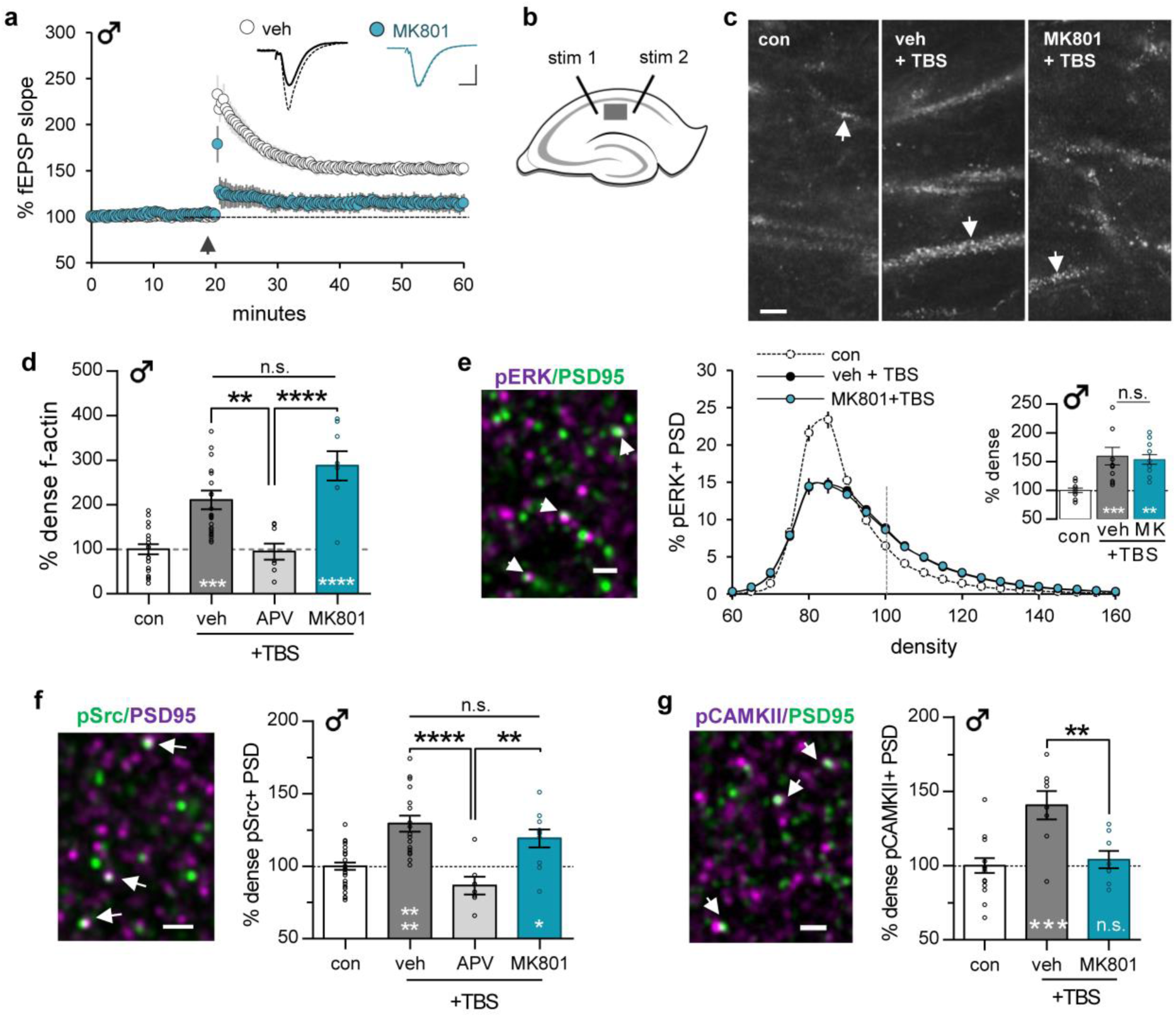
Theta burst stimulation (TBS) elicits non-ionic NMDAR signaling and actin polymerization. Stimulation was applied to Schaffer-commissural (SC) projections in slices from adult male rats. (**a**) TBS (arrow) elicits robust CA1 LTP in vehicle (veh)-treated slices whereas MK801 (30 µM, introduced 2 hr before TBS) blocked this effect (veh N=8, MK801 N=5). Traces from before (solid) and 40 min after (dashed) TBS. (**b**) For C-G, stimulating (stim) electrodes were placed in CA1a and c; analysis focused on CA1b stratum radiatum (gray box). (**c**) Phalloidin labeling in slices receiving low-frequency control (con) stimulation or TBS in the presence of veh or MK801 (30µM); arrows indicate phalloidin-labeled elements. (**d**) TBS increased numbers of densely phalloidin-labeled spines above measures after control stimulation; this effect was blocked by APV (100µM) but not MK801 (F_3,57_=15.30, p<0.0001; N=8-24). (**e-g**) Fluorescence deconvolution tomography was used to access NMDAR contributions to TBS-induced signaling. (**e**) Deconvolved images show pERK- and PSD95-immunoreactivity (ir); double-labeling appears white (arrowheads). TBS caused a rightward-skew (towards greater densities) in the density-frequency distribution for synaptic pERK-ir (F_38,608_= 18.50, p<0.0001; p=0.0048 post-hoc); this was unaffected by MK801. Inset: mean numbers of densely pERK-immunoreactive spines (≥100 density units) normalized to control slice values (F_2,32_=10.33, p=0.0003; N=11-12/group). (**f**) TBS-induced increase in numbers of synapses with dense pSrc (Y419)-ir was blocked by APV (100µM) but not MK801 (F_3,62_=15.11, p<0.0001; N=7-32/group). (**g**) TBS increased synaptic pCaMKII-ir and this effect was blocked by MK801 (F_2,29_=10.53, p=0.0004; N=8-16/group). Bars: (**a**) 1mV, 10ms; (**c**) 5 µm; (**e-g**) 2 µm. Statistics: (**a**) two-tailed unpaired t-test; (**e**) two-way repeated-measures ANOVA (interaction) with Bonferroni post-hoc tests; (**d, e** inset**, f, g**) One-way ANOVA with Tukey post-hoc tests. Asterisks inside bars denote significance vs. con stimulation. Asterisks above bars denote significance between TBS groups. n.s. = not significant, *p<0.05, **p<0.01, ***p<0.001, ****p<0.0001. Mean ± s.e.m. values shown. **Table S1** contains detailed statistics.

Next, we tested the MK801 sensitivity of TBS effects on phosphorylation of three NMDAR-driven kinases that play important roles in actin management and LTP. Slices were harvested within 15-minutes of TBS and processed for immunofluorescence localization of phosphorylated (p) ERK1/2, pSrc, or pCaMKII co-localized with postsynaptic marker PSD95. Fluorescence Deconvolution Tomography (FDT) was used to make 3-D reconstructions of the sample field and quantify immunolabeled profiles that fell within the size constraints of synapses. The density of immunolabeling for the phosphorylated protein at each double-labeled profile was measured and the resultant values were used to construct fluorescence intensity frequency distributions representing 80-120 thousand synapses per slice. Consistent with earlier work^22^, TBS caused a rightward skew of the frequency distribution, towards greater labeling densities, for synaptic p-ERK in vehicle-treated slices and MK801 did not attenuate this effect: the TBS+MK801 curve was nearly superimposed with that for TBS+vehicle (**Fig. 2e**). We confirmed previous reports^22,38^ that TBS similarly elevated synaptic pSrc immunoreactivity, and this effect is blocked by APV. But, as with pERK, the TBS-induced increase in synaptic pSrc was unaffected by MK801 (**Fig. 2f**). MK801 did, however, block the increase in synapses with dense pCaMKII that is normally induced by TBS (**Fig. 2g**). CaMKII is a calcium-dependent kinase that enables activity-driven transfer of AMPARs into synapses and is critical for LTP expression in both sexes^25,52^.

#### An antagonist of the GluN2B NMDAR subunit blocks actin polymerization

The long cytoplasmic tail domain (CTD) of GluN2B plays an important role in m-NMDAR signaling, synaptic plasticity and memory^53,54^. We accordingly evaluated the effects of Ro25-6981 (Ro25), a selective allosteric antagonist of GluN2B^55,56^, on actin polymerization and LTP in slices from male rats. First, we tested if Ro25 (3µM) depressed pharmacologically isolated NMDAR-mediated responses in CA1 field recordings. A cocktail composed of antagonists of AMPARs (DNQX: 20µM) and GABA_A_Rs (picrotoxin: 30µM) eliminated ∼90% of the fEPSP. Subsequent infusion of MK801 confirmed that the residual response was mediated by NMDARs (**Fig. 3a**). Ro25 did not measurably affect these NMDAR-gated fEPSPs (**Fig. 3b**). However, it reduced NMDAR-mediated EPSCs by ∼25% in clamp recordings (**Fig. 3c**). The clamp effect agrees with earlier work that also established an exclusively synaptic location of GluN2B in CA1^57^. The discrepancy between the extracellular vs. whole cell recording results likely reflects the pronounced difference in membrane depolarization generated in the two approaches and thus the degree to which the NMDAR’s voltage-sensitive magnesium block is reduced. The results also accord with suggestions that GluN2B di-heteromeric receptors – the presumed targets for Ro25 – are present at low levels in CA1 synapses relative to GluN2A di-heteromers and tri-heteromers^58^.

**Fig. 3.**
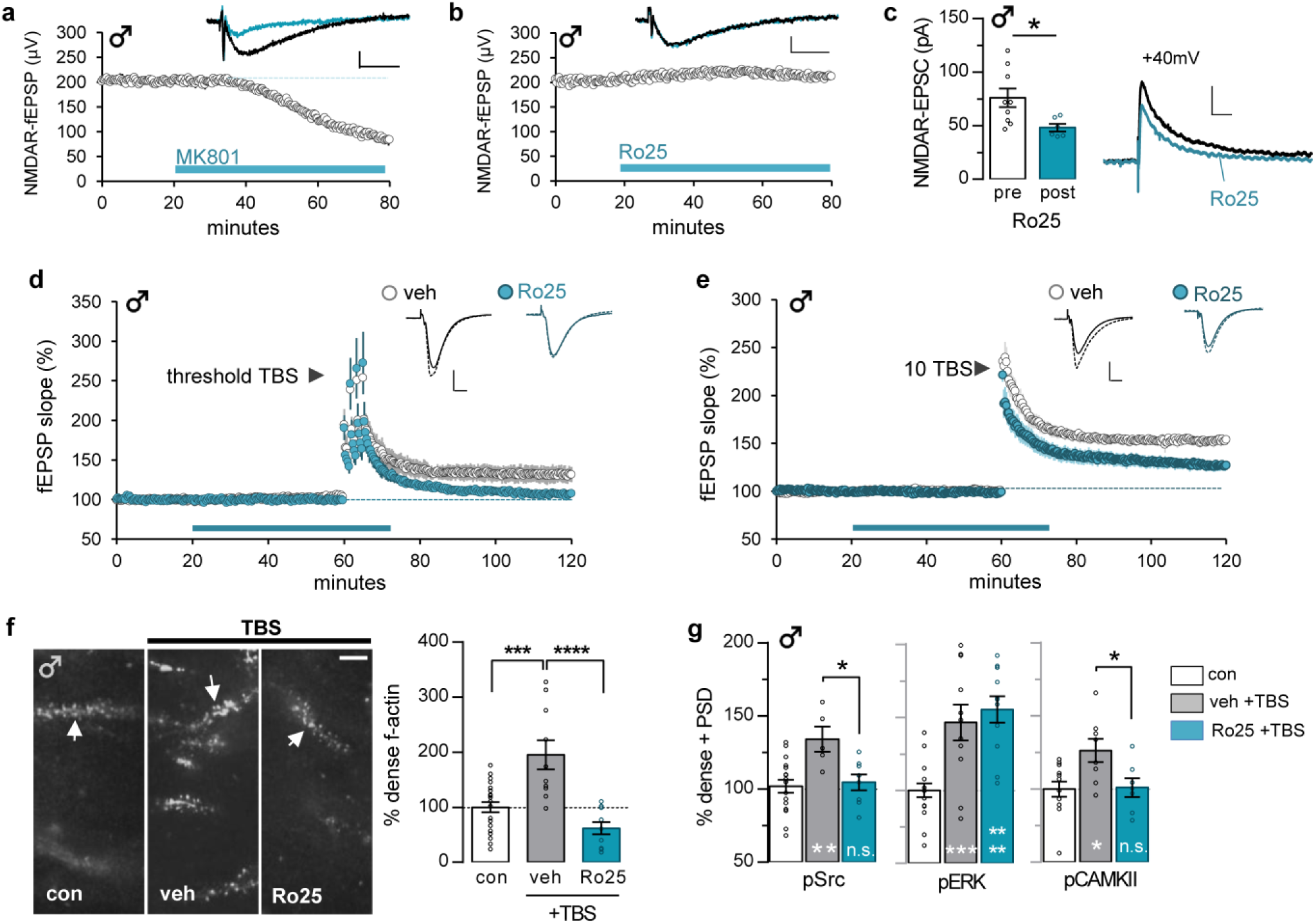
GluN2B antagonist Ro25-6981 blocks TBS-induced kinase activation, actin polymerization, and LTP in males. (**a,b**) The NMDAR-mediated component of CA1 fEPSPs was isolated using AMPAR antagonist DNQX (20µM) and GABA_A_R antagonist picrotoxin (30µM). The isolated NMDAR response was depressed by MK801 (30µM; N=6) **(a)** but not by Ro25-6981 (Ro25; 3µM; N=5) (**b**). (**c**) Voltage-clamp recordings from adult mouse CA1 pyramidal cells held at +40mV show that Ro25 infusion decreased NMDAR-EPSC amplitude (p=0.027; pre-Ro25 N=9, post-Ro25 N=6). (**d,e**) Ro25 (horizontal bar) reduced SC-CA1 LTP induced by (**d**) threshold-level TBS (4 TBS triplets, 90s intervals; N=5/group) or (**e**) a 10-burst TBS train (veh N=7, Ro25 N=9). (**f-**(**g**) Slices received either control (con) low-frequency stimulation or 10 burst TBS in the presence of vehicle (veh) or Ro25. (**f**) Phalloidin F-actin labeling (left) and labeled puncta quantification (right) show that TBS doubled the numbers of spines with dense F-actin in veh-treated slices and this effect was completely blocked by Ro25 (F_2,39_=16.81, p<0.0001, N=10-22/group, values normalized to con mean). (**g**) Ro25 blocked the TBS-induced increase in numbers of PSD95+ synapses with dense pSrc and pCaMKII but not pERK immunolabeling (pSrc: F_2,27_=6.517, p=0.0049; N=5-17/group, pERK: F_2,36_=14.36; p<0.0001, N=11-17/group; pCaMKII: F_2,24_=5.111, p=0.0142; N=7-12/group). Bars: (**a,b**) 100µV, 20ms; (**c**) 50pA, 50ms; (**d-f**) 1mV, 10ms; (**g**) 5μm. Statistics: two-tailed paired (**a-c**) and unpaired (**d,e**) t-test, (**f,g**) one-way ANOVA with post-hoc Tukey. Asterisks inside bars denote experimental vs. control comparisons; black asterisks indicate experimental group comparisons. n.s. = not significant, *p<0.05, **p<0.01, ***p<0.001, ****p<0.0001. Mean ± s.e.m values shown. **Table S1** contains detailed statistics.

Despite its minimal effects on NMDAR-mediated fEPSPs, in slices from males Ro25 impaired LTP that was induced by near threshold levels of TBS. The initial expression of potentiation was unaffected by Ro25 but responses returned to near baseline levels after an hour (**Fig. 3d**). Ro25 also reduced LTP generated by a full-length train of 10 theta bursts (**Fig. 3e**). In agreement with results summarized in Fig. 3b, the drug did not influence within-train facilitation of fEPSP responses during TBS (**Fig. S2**).

TBS-induced increases in spine F-actin were entirely blocked by Ro25 in slices from adult male rats (**Fig. 3f**), consistent with the drug’s actions on LTP consolidation. Ro25 also eliminated the effects of TBS on pSrc at CA1 synapses but did not attenuate the pERK response (**Fig. 3g**). GluN2A-containing NMDARs, which are known to upregulate ERK phosphorylation^59^ independently of calcium^60^, together with TrkB receptor signaling^61^ might explain this result. Ro25 treatment blocked TBS-driven increases in synaptic pCaMKII-immunoreactivity, indicating that both ionic and non-ionic NMDAR functions are needed to engage this LTP-critical protein.

### Females do not use GluN2B signaling for stabilization of LTP

MK801 blocked both STP and LTP in slices from females (**Fig. 4a**), but had no effect on TBS-induced increases in spine F-actin. The latter effect was eliminated by APV (**Fig. 4b,c**). However, in striking contrast to males, Ro25 did not measurably affect TBS-induced increases in spine F-actin (**Fig. 4c**) or the LTP magnitude elicited by TBS (**Fig. 4d**). Recent work showed that in females, but not males, estrogen receptor alpha (ERα) is critical for TBS-driven activation of various kinases upstream from F-actin assembly^22^. Consistent with this, the ERα antagonist MPP (3μM) prevented TBS-induced increases in spine F-actin in females, but not in males (**Fig. 4e**). This result suggests that females may substitute local estrogen signaling for m-NMDAR operations evident in males, for rapid activity-induced remodeling of actin networks in mature spines.

**Fig. 4.**
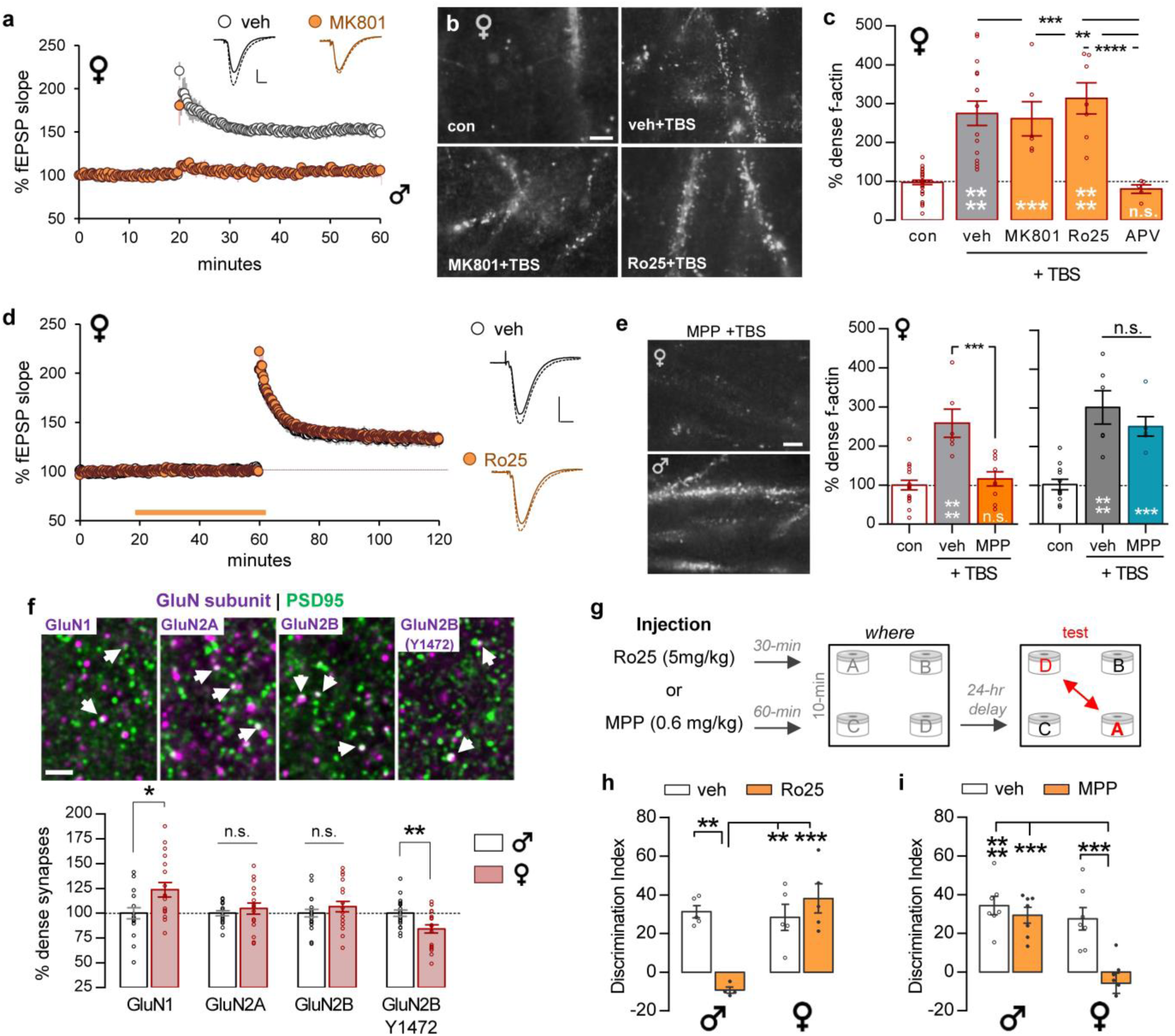
GluN2B antagonist Ro25-6981 does not block TBS-driven actin polymerization, CA1 LTP, or episodic memory in females. **a-e)** Electrode placements as in Figure 2b. (**a**) MK801 (30µM) blocked TBS-induced LTP in female rats (vehicle, veh N=5, MK801 N=4). (**b**) Phalloidin labeling in CA1 of slices that received control, low frequency SC stimulation (con) or 10 burst TBS in the presence of vehicle, MK801, or Ro25 (3µM). (**c**) TBS increased phalloidin labeled spine F-actin levels in the presence of vehicle, MK801, or Ro25 (vs. controls) but this increase was blocked by NMDAR antagonist APV (F_4,62_=22.88, p<0.0001; N=5-33, values normalized to con mean). (**d**) Ro25 did not disrupt TBS-induced LTP in female slices (veh N=5, Ro25 N=6). Traces from before (solid) and 60 min after (dashed) TBS. (**e**) In vehicle-slices, TBS increased numbers of densely phalloidin labeled puncta in both sexes. This effect was blocked by ERα antagonist MPP (3µM) in females (F_2,29_=16.02, p<0.0001; N=6-17) but not in males (F_2,21_=20.28, p<0.0001; N=6-12). (**f**) Deconvolved images of NMDAR subunit and PSD95 immunolabeling in CA1; arrows indicate double-labeled profiles. In females, the % PSD95^+^ synapses with dense GluN1-immunoreactivity (ir) was greater than in males (p=0.0171) whereas levels of GluN2A- and GluN2B-ir were comparable (N=17-20/group). The percent PSD95^+^ synapses with dense pGluN2B Y1472-ir was lower in females than males (p=0.0041; N=17-20/group). (**g**) Mice received vehicle, Ro25 or MPP before odor exposure in the 4-corner episodic ‘where’ paradigm. (**h,i**) Vehicle-treated males and females (2 cohorts) discriminated the moved cues in the episodic ‘where’ task; (**h**) Ro25 disrupted this effect in males (veh N=5, Ro25 N=4) and had no effect on female performance (F_1,15_=19.62, p=0.0005; female N=5/group). (**i**) In contrast, MPP blocked this ‘where’ acquisition in females but did not attenuate performance in males (F_1,24_=8.001, p=0.0093; N=7/group). Scale bar: (**a,d**) 1mV, 10ms; (**b,e**) 5μm; (**f**) 2μm. Statistics: two-tailed unpaired t-test (**a,d**), (**f**) two-tailed unpaired t-test Welch’s correction, (**c,e**) one-way ANOVA with Tukey post-hoc, (**h,i**) 2-way ANOVA with post-hoc Tukey. Asterisks inside bars denote comparison to controls; n.s. = not significant, *p<0.05, **p<0.01, ***p<0.001, ****p<0.00001. Mean ± s.e.m values shown. **Table S1** contains detailed statistics.

The failure of Ro25 to disrupt actin polymerization and LTP in females raises the possibility of sex differences in concentrations or post-translational modifications of synaptic GluN2B subunits. FDT analysis showed a modest sex difference in synaptic concentrations of GluN1 but not GluN2A or GluN2B (**Fig. 4f**). However, synaptic GluN2B Y1472 phosphorylation^62^ was significantly lower in females than males (p=0.0041, 2-tailed unpaired t-test), suggesting a plausible explanation for the more prominent role of GluN2B in LTP stabilization in males than females.

### Sex differences in metabotropic signaling underlying memory

Together the aggregate LTP results and the expectation that CA1 LTP is critical for learning in the episodic tasks, give rise to the prediction that blocking GluN2B-mediated m-NMDAR signaling will more severely impair episodic learning in males than in females and, conversely, that blocking ERα will disrupt this learning in females but not males. We tested this by treating mice with vehicle, Ro25 (5mg/kg, 30 min), or MPP (0.6 mg/kg, 60 min) prior to initial odor exposure in the simultaneous cue ‘where’ task^5,21^ (**Fig. 4g**). In this paradigm, both sexes showed positive retention scores when given 10-min training and tested 24 hours later. Ro25 produced a profound deficit for encoding ‘where’ information by males without effect in females (**Fig. 4h**). Conversely, the ERα antagonist MPP eliminated discrimination of the moved cues in females without attenuation of learning in males (**Fig. 4i**). Analyses of total sampling times and locomotor activity across sex and trials showed that the effects of Ro25 and MPP on behavior were not due to a reduction in arousal or general interest in the cues (**Fig. S3**). Thus, males and females rely upon non-ionic signaling from different types of receptors for encoding at least one component of episodic memory.

## Discussion

As is the case with memory, LTP passes through several consolidation stages during which it is vulnerable to disruption^63^. The underlying processes involve multiple small GTPase-initiated signaling cascades that lead to formation and subsequent stabilization of actin filaments^40^. These events serve to anchor a change in the shape of the dendritic spine and its postsynaptic specialization^24,63^. The present studies led to the surprising conclusion that this complex collection of events, while dependent on NMDARs, can be completed without calcium flux through those receptors. Thus, in both males and females, blocking the receptor channel entirely eliminated LTP but left intact TBS-induced actin polymerization (‘consolidation without potentiation’).

There were however important differences in the manner in which the two sexes executed non-ionic, actin regulatory signaling as evidenced by the male-specific effects of the GluN2B antagonist Ro25-6981. The long GluN2B-CTD associates with SynGAP, which controls activity of the small GTPase Ras and thereby regulates the actin severing protein cofilin and actin polymerization^64^. SynGAP also potently influences the activity of Rap^65^, a GTPase intimately involved in integrin activation^66^. Integrins regulate the actin cytoskeleton at various adhesion junctions and are essential for initiating TBS-induced F-actin assembly in hippocampus^45,67^. Although females rely on the NMDAR, it appears that they substitute local release of estrogen onto ERα for the GluN2B-dependent m-NMDAR actions that are critical for actin polymerization and LTP in males. Thus, both sexes use a combination of ionic and metabotropic operations to modify synapses but execute the latter function in radically different ways.

Synaptic ERα levels are substantially greater in females than males and estradiol acts through ERα to activate postsynaptic Src and ERK in females only^22^. These findings help explain why male rodents, despite having high estrogen levels in hippocampus^68^, do not use the hormone to promote LTP. Why the male m-NMDAR mechanism is missing in females is not known but we found the synaptic GluN2B Y1472 site to be more intensely phosphorylated in males. This NMDAR CTD residue is targeted by Src family kinases, which are known to up-regulate NMDAR function^69^. Evidence that estrogen decreases phosphorylation of GluN2B Y1472^70^ raises the possibility that the same estrogen signaling needed for consolidation of female LTP suppresses the metabotropic NMDAR activities engaged in males (**Fig. 5**).

**Fig. 5.**
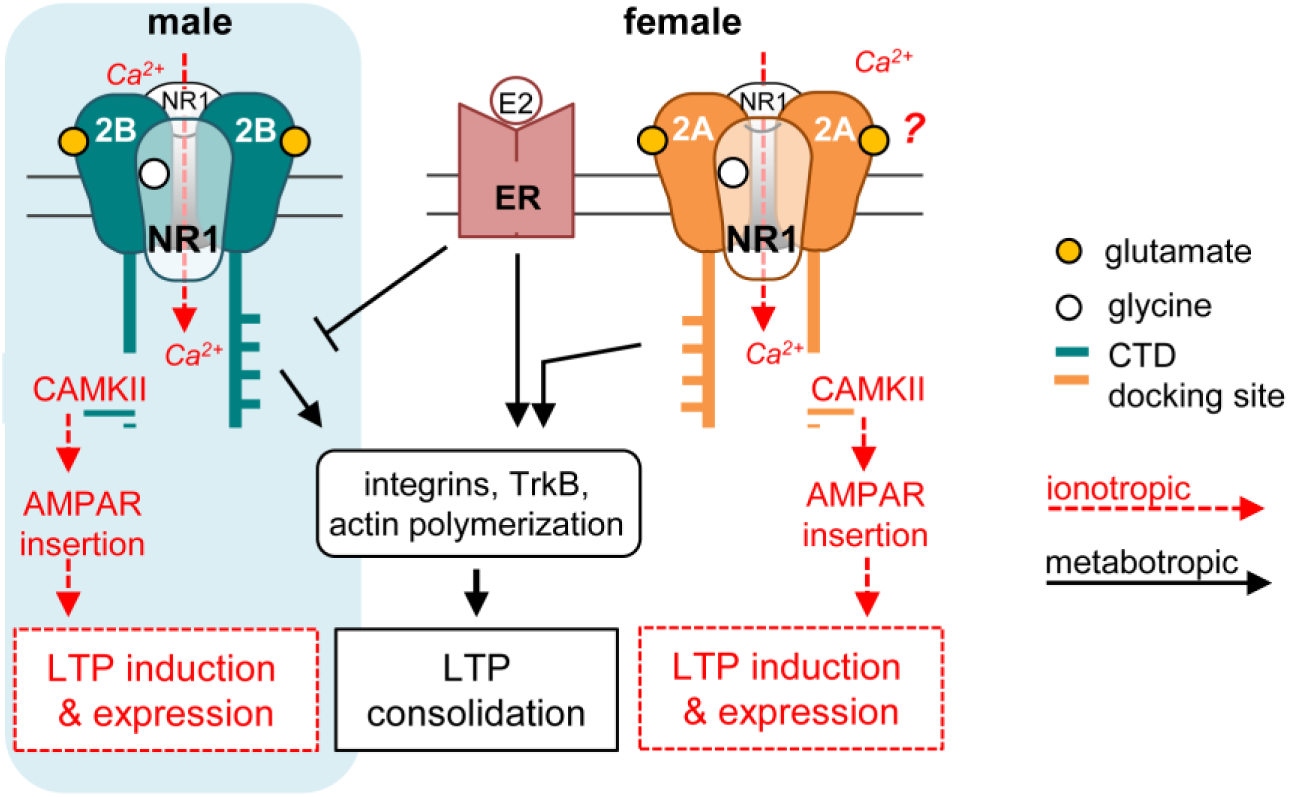
Schematic illustration of sex differences in the contributions of non-ionic NMDAR signaling to memory-related synaptic plasticity. Observed effects of APV and MK801 indicate that both sexes use ionotropic NMDAR functions to activate CaMKII and associated processes (AMPAR insertion) required for the induction of LTP. Both sexes also use non-ionic (APV-but not MK801-sensitive) NMDAR functions to trigger actin polymerization and LTP consolidation. Actions of Ro25-6981 indicate GluN2B subserves these non-ionic NMDAR functions in males, presumably via its cytoplasmic terminal domain (CTD). Females do not use the GluN2B mechanism for stable LTP; rather they rely upon activation of synaptic estrogen receptors (‘ER’) to engage the same effectors as used by males. We hypothesize that the ERs tonically suppress (male) GluN2B activities by reducing phosphorylation of the CTD Y1472 site. As noted, the APV / MK801 results suggest that females use some type of non-ionic NMDAR signaling to engage LTP consolidation machinery. One possibility illustrated is that the GluN2A CTD serves this role.

The different types of metabotropic signaling were associated with striking sex differences in episodic memory: males had a strong advantage on encoding the ‘where’ component whereas females had better scores on ‘what’ and when’. Female mice were similarly able to encode a longer cue sequence than males. Although the female results are unprecedented for rodent work, they do have correspondences in human studies, including the observation that woman outperform men when dealing with extended lists^71-73^. Critically, the GluN2B antagonist that blocked actin signaling and LTP in males but not females, had corresponding sex-specific effects on encoding the ‘where’ component of episodic memory. In this same paradigm, blocking ERα disrupted learning in females only, paralleling the female-specific effects of the ERα antagonist on LTP and the downstream signaling^22^ regulating actin polymerization.

The difference in consolidation mechanisms provides a reasonable explanation for the higher female threshold for production of stable LTP described in previous studies^22^. Specifically, in females the added need to generate the locally-derived estrogen^23,74,75^ and activate synaptic estrogen receptors likely increases the activity threshold for enduring plasticity. The relative advantages and disadvantages of a higher threshold for encoding elements of episode would relate naturally to the present behavioral results. For example, a higher plasticity threshold might be seen as a disadvantage in that it could limit the encoding of weaker cues but this same constraint might be an advantage for preferentially encoding salient cues relative to background elements of lesser significance.

Questions inevitably arise about the extent to which sex differences in human learning reflect societal perceptions and expectations along with educational practices^15,16^. While these factors undoubtedly contribute in people, our results show that the female advantage for encoding non-spatial aspects of episodic memory is present when such considerations are absent, as was male advantage in earlier tests of episodic ‘where’ acquisition^21^. Moreover, the differences in facility for acquiring different components of episodic memory are associated with dramatic sex differences in the synaptic machinery for encoding. We therefore conclude that sex differences in episodic memory have biological as well as social origins.

## Author Contributions

Conceptualization: AAL, JCL, CMG, and GL

Methodology: AAL, JCL, CMG, and GL

Investigation: AAL, JCL, YJ

Software: CDC

Writing – original draft: AAL, JCL, CMG, and GL

Writing – review & editing: AAL, JCL, CMG, and GL

## Materials & Correspondence

The data supporting the findings of this study are available from the corresponding author upon reasonable request. Theta burst areas for field electrophysiology were analyzed using code available at https://github.com/cdcox/Theta-burst-analyzer-for-Le-et-al. Code for FDT analysis is made available upon request. The use of the FDT code is strictly prohibited without a Licensing Agreement from The University of California, Irvine.

## Supplementary Figures for “Metabotropic NMDA Receptor Signaling Contributes to Sex Differences in Synaptic Plasticity and Episodic Memory”

**Fig. S1.**
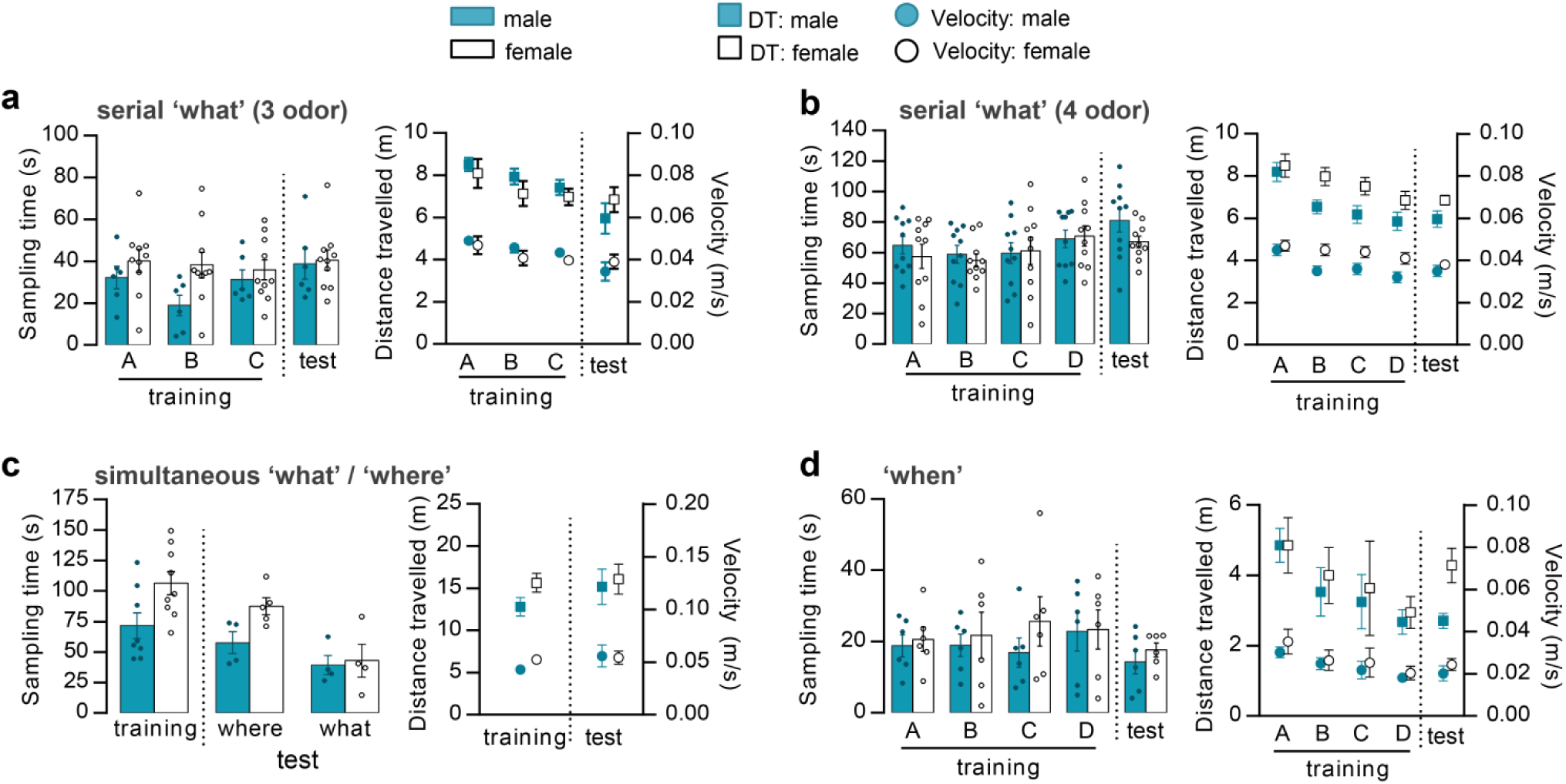
Detailed locomotor activity and sampling times in the behavioral tasks. (**a**) Serial 3-odor ‘what’ task (presented in Figure 1a). *Left*: The time spent sampling odors in the serial training (A-B-C) and test trials was similar across groups (interaction: p>0.05). *Right*: Distance traveled (DT, squares) and movement velocity (circles) during each trial were also similar (interaction: p>0.05). (**b**) Sampling and locomotor data for the Serial 4-odor ‘what’ task (Fig 1A) (interaction: p>0.05). (**c**) Sampling times and locomotor data during training for simultaneous ‘what’ and ‘where’ (from Fig 1b,d) were pooled together due to having the same initial (training) trial (interaction: p>0.05). (**d**) Sampling and locomotor activity for the ‘when’ task (from Fig 1c) (interaction: p>0.05). Statistics were performed with two-way ANOVA. For all panels, N=4-10/group. Mean and s.e.m. shown. **Table S1** contains detailed statistics.

**Fig. S2.**
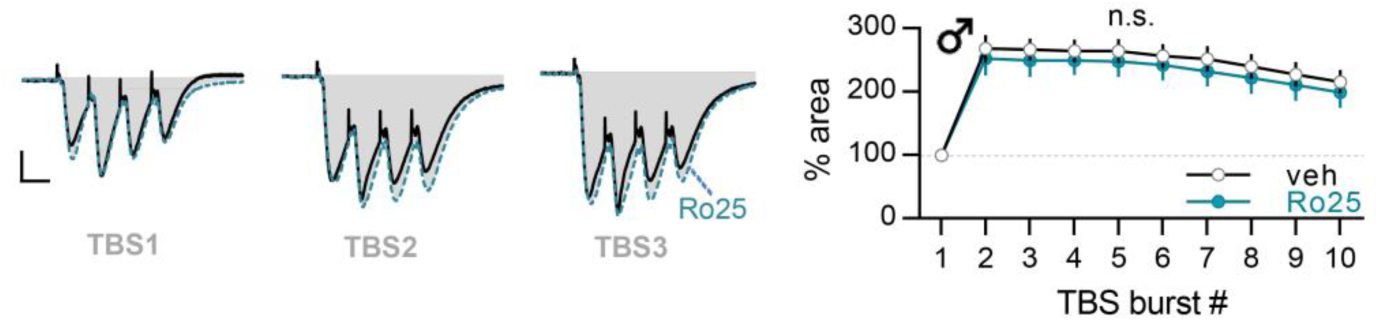
GluN2B antagonism did not reduce the size of fEPSP responses with TBS. Vehicle (veh) or Ro25-6981 (Ro25, 3µM) was infused into acute male rat slices for 40 minutes, and then 10 burst TBS was delivered to the CA3-CA1 projections. The area of the response to each burst was normalized to that of the first burst (TBS1) response. Traces at left show representative fEPSP response areas (shaded gray) to the first three bursts of the 10 theta burst train in veh- (black) and Ro25-6981- (Ro25; blue, dashed) treated slices. The line graph shows group mean values for each pulse in the 10 burst train. The response areas were unaffected with Ro25 treatment (F_9,189_=0.2407, p=0.9880, two-way repeated measures ANOVA. N=15 veh, N=9 Ro25). Scale bar: 1mV, 10ms. Mean and s.e.m. shown. **Table S1** contains detailed statistics.

**Fig. S3.**
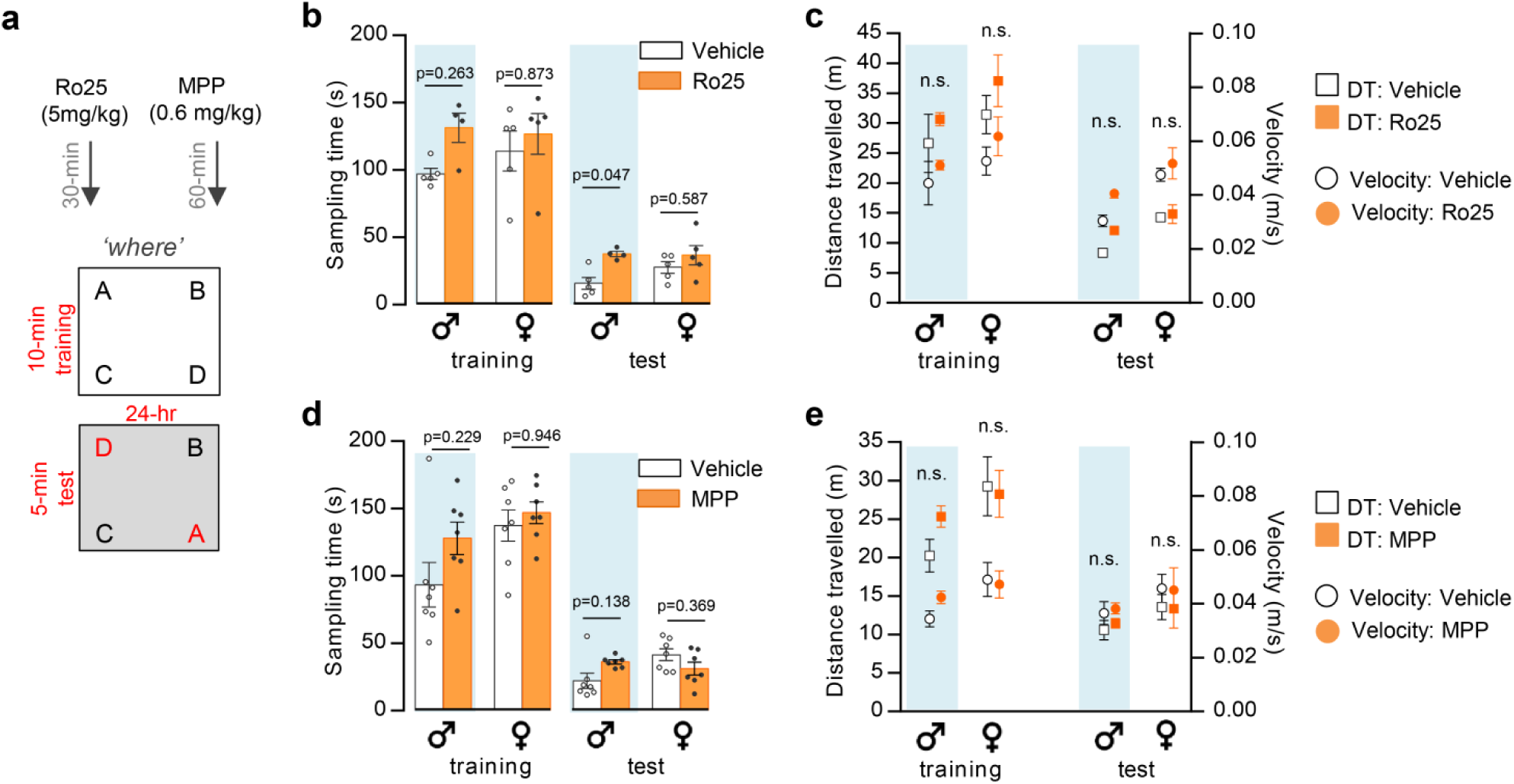
Locomotor activity and sampling times in the “where” task were not reduced by Ro25-6981 (Ro25) or MPP. (**a**) Schematic detailing the protocols (retention data is presented in Fig 4h for Ro25 and Fig 4i for MPP). (**b-e**) Graphs show that the sampling times (b,d), and the velocity and distance traveled (c,e), for all drug-treated groups were not lower than their respective vehicle-treated groups; the Ro25 drug was found to increase sampling times in males during the test phase (b) but other measures were not affected (p-values at top of columns are for vehicle vs. drug post-hoc comparisons for each measure; n.s., not significant, p>0.05). In addition, across all measures there was no statistical difference for effect of drug between sexes during either training or testing. For both training and test phases, data were analyzed by 2-way ANOVA (sex and drug) followed by Tukey post hoc comparisons; See **Table S1** for detailed statistics. Mean and s.e.m. shown.

**Supplementary Table 1:**

- *see attached .xlsx file*.

